# Optimal Scheduling of Bevacizumab and Pemetrexed/cisplatin Dosing in Non-Small Cell Lung Cancer

**DOI:** 10.1101/540849

**Authors:** Benjamin K Schneider, Arnaud Boyer, Joseph Ciccolini, Fabrice Barlesi, Kenneth Wang, Sebastien Benzekry, Jonathan P Mochel

## Abstract

Bevacizumab-pemetrexed/cisplatin (BEV-PEM/CIS) is a first line therapeutic for advanced non-squamous non-small cell lung cancer (NSCLC). Bevacizumab potentiates PEM/CIS cytotoxicity by inducing transient tumor vasculature normalization. BEV-PEM/CIS has a narrow therapeutic window. Therefore, it is an attractive target for administration schedule optimization. The present study leverages our previous work on BEV-PEM/CIS pharmacodynamic modeling in NSCLC-bearing mice to estimate the optimal gap in the scheduling of sequential BEV-PEM/CIS. We predicted the optimal gap in BEV-PEM/CIS dosing to be 2.0 days in mice and 1.2 days in humans. Our simulations suggest that the efficacy loss in scheduling BEV-PEM/CIS at too great of a gap is much less than the efficacy loss in scheduling BEV-PEM/CIS at too short of a gap.

## Introduction

Bevacizumab-pemetrexed/cisplatin (BEV-PEM/CIS) combination therapy has been shown to be an effective first line and maintenance therapy for non-small cell lung cancer (NSCLC) in Phase II and Phase III clinical trials(1–3). Pemetrexed inhibits enzymes necessary for pyrimidine and purine synthesis – primarily thymidylate synthase, which is necessary for thymidine synthesis and tumor cell replication(4). Cisplatin is an alkylating agent which crosslinks adjacent N7 centers on purine residues, damaging DNA, disrupting repair, and disrupting purine synthesis(5–7). Disrupting DNA substrate supply results in S-phase arrest, DNA repair disruption, and eventually apoptosis(8,9). Cisplatin also significantly disrupts calcium and reactive oxygen species (ROS) regulation, inducing cellular lesions which further sensitizes cancer cells to apoptosis(7).

In contrast to the effect of PEM/CIS, i.e. DNA damage, bevacizumab is an anti-VEGF (vascular endothelial growth factor) humanized monoclonal antibody. VEGF is an angiogenic potentiator which promotes the growth of endothelial tissue necessary for arteries, veins, and lymphatics. By limiting neovascular growth, and therefore blood delivery to neoplasms, bevacizumab exhibits limited antiproliferative properties(10).

More importantly, bevacizumab transiently induces a pruning effect on neovascular beds, which normalizes blood supply to neovascularly dense tissues (i.e. tumors)(11–13). By normalizing blood supply, bevacizumab enhances chemotherapeutic (i.e. PEM/CIS) delivery to neoplasms(14,15).

The effects of BEV-PEM/CIS are generalized i.e. any cell capable of uptaking the drugs are susceptible to their effects, especially rapidly-dividing cells such as myeloid cells(16). Accordingly, BEV-PEM/CIS has a narrow therapeutic window and generalized side-effects(3). Previous studies on BEV-PEM/CIS suggest that sequential administration of BEV-PEM/CIS (i.e. BEV before PEM/CIS) outperforms concomitant scheduling of BEV-PEM/CIS in treating NSCLC(12,17–19). This makes BEV-PEM/CIS an attractive target for scheduling optimization via modeling and simulation, as a range of practical predictions – such as optimal scheduling in humans – can be made without the considerable time and resource investment required to conduct *in vivo* experiments. These predictions can be used to guide future studies, greatly accelerating drug development(20).

In our previous work on BEV-PEM/CIS published in Imbs et al. 2017(17), mice with NSCLC tumor xenografts were administered bevacizumab with PEM/CIS combination therapy (CT) in either concomitant, or delayed (i.e. bevacizumab before PEM/CIS) scheduling. The NSCLC tumors had been modified such that tumor growth could be tracked over time via either *bioluminescence* or *fluorescence.* Following previous theoretical investigations, the dataset generated from the mice with *bioluminescent* tumors was used to develop a semi-mechanistic PK/PD model for tumor dynamics in response to BEV-PEM/CIS(21,22). The model was then used to predict the optimal scheduling gap between bevacizumab and PEM/CIS administration.

The aim of this follow-up modeling work was to both refine and expand upon previous results on BEV-PEM/CIS CT using the much larger *fluorescence* dataset generated in Imbs et al. 2017(17). We first showed that the semi-mechanistic model previously developed better explained the data than comparable models (i.e. we validated the previously developed structural model). Then, we refined the parameter estimates of the model, and used it to predict the optimal scheduling gap between bevacizumab and PEM/CIS administration. Next, we used stochastic simulations to explore the marginal loss in therapeutic efficacy when BEV-PEM/CIS was administered at a sub-optimal gap, the effect of bevacizumab dose scaling on population optimal gap, as well as the inter-individual variability (IIV) of optimal gap. Lastly, using literature human PK/PD models and parameter estimates, we were able to scale the model to estimate the optimal scheduling of BEV-PEM/CIS in humans.

## Methods

### Experimental Procedure

Comprehensive details on animals and the experimental procedure are available in Imbs et al. 2017(17). Briefly, on Day 0 of the experiment, tumors (ca 120,000 cells) consisting of H460 human NSCLC transfected with luciferase and the tdTomato gene (H460 Luc+ tdTomato+, Perkin Elmer France) were injected ectopically into the left flank of 90 mice. Animals were pathogen-free, immunocompromised, 6-week-old, female Swiss nude mice (Charles River, France). The mice were randomized into one of five treatment groups. The first study group (Control) received no treatment. The second treatment group (PEM/CIS) was administered both 100 mg/kg of pemetrexed IP and 3 mg/kg of cisplatin IP on Days 14, 28, and 42 of the experiment. The third, fourth, and fifth treatment groups (BEV-PEM/CIS) received the same PEM/CIS treatment as the second experimental group.

In addition, the BEV-PEM/CIS treatment groups were administered 20 mg/kg IP of bevacizumab either concomitantly with the PEM/CIS administrations (Group 3), 3 days prior to each PEM/CIS administration (Group 4), or 8 days prior to each PEM/CIS administration (Group 5) – see Table S1 for administration tabulation.

Tumor growth was monitored on a minimum bi-weekly basis using Ivis Spectrum imager (Perkin Elmer France) and images were acquired and analyzed using the Living Image 6.0. software (Perkin Elmer France).

Mice were supplied with paracetamol supplemented water (e.g. 80 mg/kg/day) to prevent disease-related pain. Animals showing signs of distress, pain, cachexia (i.e. loss of 10% of body mass), or extensive tumor proliferation (i.e. within 2-3 cm) were euthanized. All animals were euthanized on Day 87 of the experiment. All experiments were approved by the local ethical committee at French Ministère de I’Education Nationale, de I’Enseignement Supérieur et de la Recherche, and registered as #2015110616255292.

### PK/PD Structural Model Building and Evaluation

The PK models for bevacizumab, pemetrexed, and cisplatin were derived from previously published PK models in mice(23–25). The parameters for these models were fixed to the typical values from those studies and assumed no IIV.

The PD model was selected from a series of sequentially fit tumor growth and drug effect models. First – using only the control dataset – the exponential, linear-exponential, and Gompertz growth model were cross-evaluated as models of unperturbed tumor growth(26). Then, incorporating the full dataset into the fit, the log-kill effect of bevacizumab, log-kill effect of pemetrexed, and log-kill effect of cisplatin were each considered. The interaction effect between bevacizumab and PEM/CIS was included to represent the synergistic effect of bevacizumab. Following previous work for the effect of cytotoxic drugs, three cellular death compartments were included in the PD portion of the model to represent the delay between cellular damage due to PEM/CIS and cell death(27).

Competing models were evaluated numerically using Bayesian information criteria (BIC) and the precision of parameter estimates – defined as the relative standard error of the estimate (RSE).

Observed vs. predicted plots, individual fit plots, and Visual Predictive Checks (VPCs) were produced to graphically assist model evaluation (as automated in Monolix 2018R2). VPCs were produced using the default estimation process for VPCs as of Monolix 2018R2 i.e. to create the 90% prediction intervals for the 10^th^, 50^th^, and 90^th^ percentiles, 500 simulations are performed using random individual parameters and the design structure of the experiment.

After model selection, the statistical correlations between random-effects were explored via visual inspection. Correlations plots between random effects were produced, defined as vs *ηi,t,φ*_2_ i.e. the random effect *η,* of individual *i,* at time *t,* of parameter 1, i.e. *φ*_1_, vs the random effect *η*, of individual *i,* at time *t*, of parameter 2, i.e. *φ*_2_. The full posterior distribution of the parameters were used in place of EBEs to avoid visual artifacts due to shrinkage as suggested by Lavielle, et al, 2016(28) and Pelligand L, et al, 2016(29). Statistical correlations between random effects were also numerically assessed using a Pearson correlation test at a *P* < .05 threshold.

SAEM convergence and final model parameterization were graphically assessed by inspection of search stability, distribution of the individual parameters, distribution of the random effects, individual prediction vs. observation, individual fits, distributions of the weighted residuals, as well as VPCs.

The precision of parameter estimates was numerically assessed using RSE. The normality of random effects distributions, the normality of individual parameter distributions, and the normality of the distribution of residuals were each numerically assessed using a Shapiro-Wilk test (*P* < .05). The centering of the distribution of residuals (i.e. centered on 0) was numerically assessed using a Van Der Waerden test (*P* < .05).

Parameter stability was assessed by comparing parameterizations resulting from random-initial starting value selection – as implemented in the Monolix *assessment suite.* The *assessment suite* performs 5 SAEM parameterizations in series using random initial parameter values uniformly drawn from the interval from approximately 60% to 160% of final parameter estimates. The SAEM of the individual parameterizations was then tracked between runs – giving a range of parameter value estimates, RSEs, and log-likelihoods to compare. This assessment was used to ensure that the algorithm did not converge to a log-likelihood local minimum during the process of producing final parameter estimates. The settings described are the default as of Monolix 2018R2.

Simulations

Simulations were performed in R 3.4.4 using Simulx 3.3.0(30) to simulate from Monolix run files. First, a function was built which accepted treatment schedule, parameter substitutions, dose, and number of individuals as input and produced a simulated population as an output.

This function was simply a convenience wrapper of Simulx for automation purposes and was verified by reproducing VPCs per treatment group. **Simulation set 1** was used to predict the optimal gap between administration of bevacizumab and PEM/CIS. **Simulation set 2** produced an estimate of the IIV of the optimal gap. **Simulation set 3** examined the anticipated effect of varying the dose of bevacizumab on the optimal gap. **Simulation set 4** scaled predictions of BEV-PEM/CIS efficacy to humans. All simulations were of population level response (i.e. simulated without RSE or IIV) except for **simulation set 2** (simulated without RSE and with IIV) which used 1000 monte carlo samples. Further details are provided in the **supplementary methods**.

### Quality assessment

All mlxtran and R codes were assessed for quality control by an independent evaluator.

## Results

### Error Model

Measurement error was best described using a log-normal constant-error model (equation 1). The natural-log of each individual measurement, *ln*(*y_ij_*) with individual *i* and repetition *j*, was modeled as a measurement centered on the natural-log mean of *y_ij_*, over j, i.e. 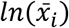, in addition to some residual error, *ϵ_ij_*, normally distributed, centered on zero, and with standard deviation *a*.

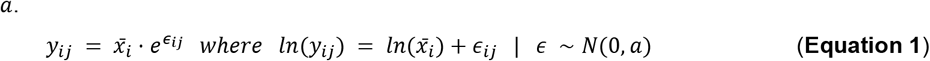

The sGOF graphics and numerical analyses supported that a log-constant error model best fit the data. The log-constant error model was not rejected for both the Kolmogorov-Smirnov test for normality (*P* = .22) nor the Pearson chi-square normality test *(P* = .0509). In contrast, a constant error model was rejected by these two tests (see **Figure S1** for further details).

### PK/PD Structural Model Building

No outliers were identified during initial data exploration. Therefore, no collected data were excluded from model building.

The pharmacokinetics of bevacizumab, pemetrexed, and cisplatin were each modeled using one-compartment models with first order IP absorption and first order elimination based on literature descriptions and PK parameter estimates(23–25). Random effects (i.e. *η_pk_*) were set to 0 as individual PK was not reported in these experiments.

A Gompertz function (**Equation 2**) was found to best describe the unperturbed tumor growth *V*(*t*), based on its fit performance over competing models, low RSE on parameter estimates, and literature-established descriptive quality.

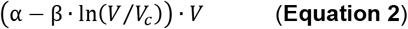

Due to sparseness in sampling, more complex semi-mechanistic models of growth were not supported by the data. Parameters *α* and *β* represent the proliferation rate of the tumor cells and rate of exponential decrease of the tumor relative growth rate, respectively. *V_c_* is the unit value of relative fluorescence units (RFU) corresponding to one cell, i.e. the proportionality constant between RFU and the number of cancer cells in the fluorescent volume.

*V_c_* was estimated externally from Monolix by conducting a naive-pooled, linear-regression on the natural-log of the full dataset. The regression gave a rough estimate of *V*_0_ which was then scaled by the approximate number of cells injected at time 0 (ca. 120,000 cells) to derive *V_c_* = 5.064 × 10^-4^ RFU.

After selecting an appropriate growth model, the log-kill effects of pemetrexed, cisplatin, and bevacizumab were each considered in parallel. The log-kill effect of bevacizumab was estimated as insignificant and removed from the model. The estimation of the log-kill effect of pemetrexed and cisplatin were found to be highly correlated. To reduce model complexity, only their combined concentration, *C*(*t*), and a corresponding log-kill parameter, *γ,* were considered in the final model (**Equation 3**).

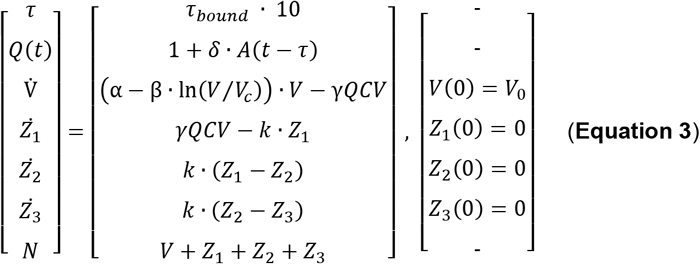

*A*(*t*) represents the plasma concentration of bevacizumab. *Q*(*t*) represents the synergistic effect of improved vascular quality. In brief, the increase in neoplasm vascular quality due to bevacizumab typically occurs within a period of a few days after administration. To represent this delay in effect, time (*t*) was delayed by *τ*. Parameter *δ* represents the proportional increase in PEM/CIS efficacy due to vascular quality improvement under bevacizumab therapy.

The estimation of *τ* was bounded between 0 and 10 using the link function *τ* = *τ_bound_* · 10, where *logit*(*τ_bound_*) ~ *N*(0,1). All other parameters were best estimated as log-normally distributed. The full statistical representation of individual parameters, *ϕ_i_*, estimated via SAEM is shown in **Equation 4**, where the full structural model is denoted by *F.*

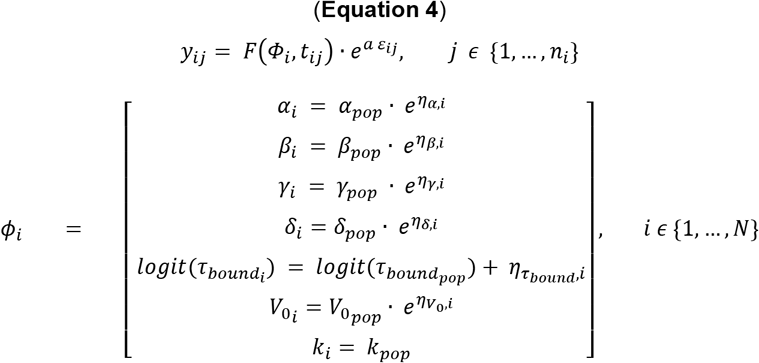

Cellular death due to chemotherapeutic treatment was modeled as a three-compartment transition from the growth compartment to death(27). The compartments are labeled *Z*_1_, *Z*_2_, and *Z*_3_ (numbering respective to their order) and transition between compartments is governed by intercompartmental clearance parameter *k*. *k* was not identifiable using the SAEM algorithm. Thus, after a period of manual exploration, *k* was set to the value of 0.3. This choice is consistent with the parameterization made in Imbs et al(17). This choice also limits the total transition time from the tumor mass compartment to cellular death to the order of a day which is consistent with upper limits of cellular death clearance(31).

The full tumor size, *N*, was the sum of the size of unperturbed cells, *V*, as well as the size of damaged cells undergoing cellular death i.e. *Z*_1_ + *Z*_2_ + *Z*_3_.

No correlations between random effects were statistically significant enough to be included in the final model (**Figure S2**). Full parameter estimates and model diagram are provided in **Table 1** and **Figure 1** respectively. Model diagnostics are collected in **Figure 2** through **Figure 4**.

**Figure 1.**
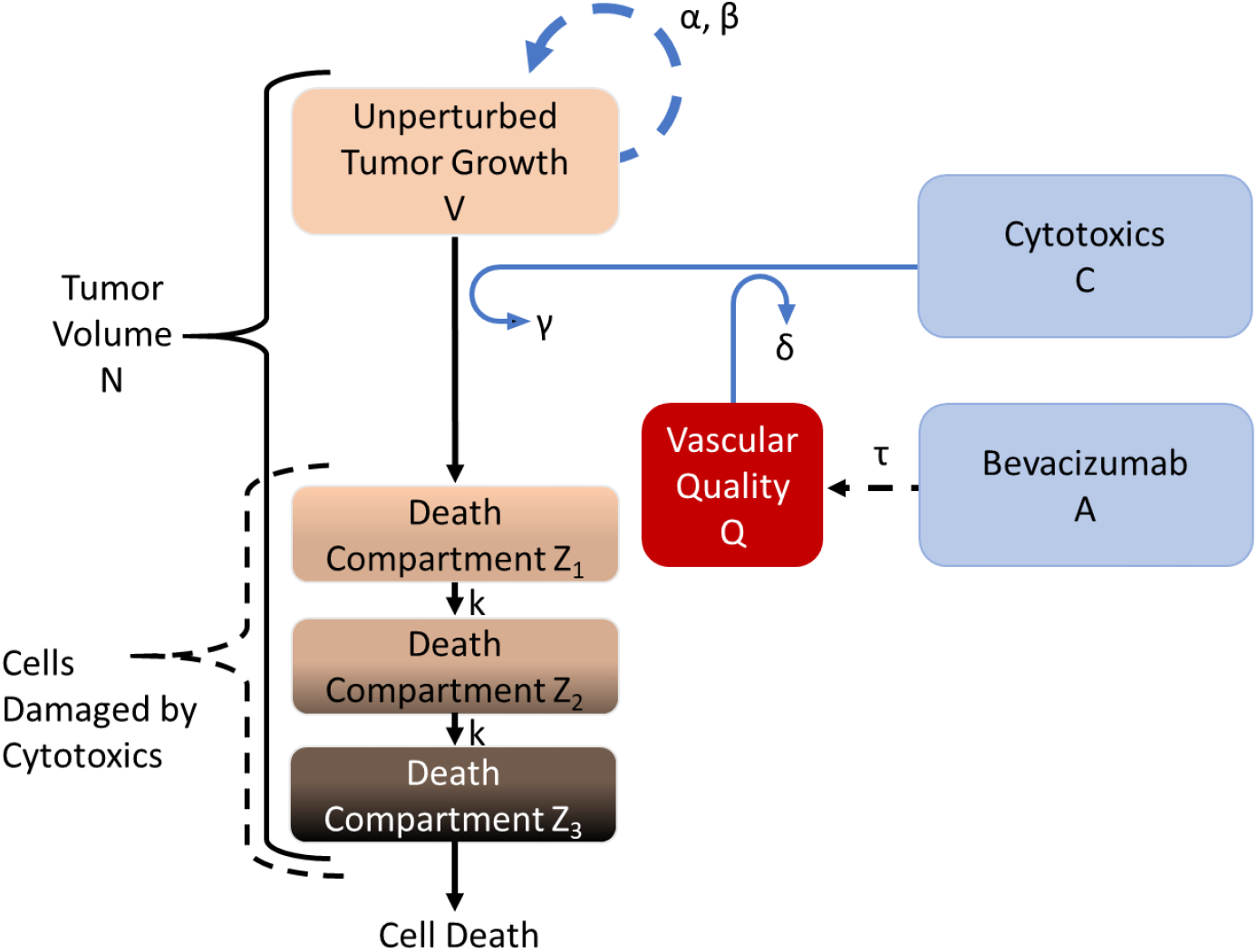
Structural Model Diagram. The scheme of the structural model is depicted to the right. Unperturbed cells grow at rate governed by *α* and *β*. When a cytotoxic is introduced into the system, the cytotoxic impairs the growth of the tumor by sending cells into a death succession. The parameter which determines the cytotoxic efficacy, *y*, is scaled by both the concentration of cytotoxics, *C*(*t*), and the volume of the tumor, *V*(*t*). Bevacizumab improves vascular quality, *Q*(*t*), after time delay, *τ*, which scales the cytotoxic effect by parameter *δ*. When a cell is damaged by cytotoxics it begins a progression from unperturbed growth – compartment *V*(*t*) – to damage compartments *Z*_1_ through *Z*_1_. Eventually the cell exits the tumor volume as it dies. The rate of transfer between damage compartments is governed by intercompartmental clearance parameter *k.*

**Figure 2.**
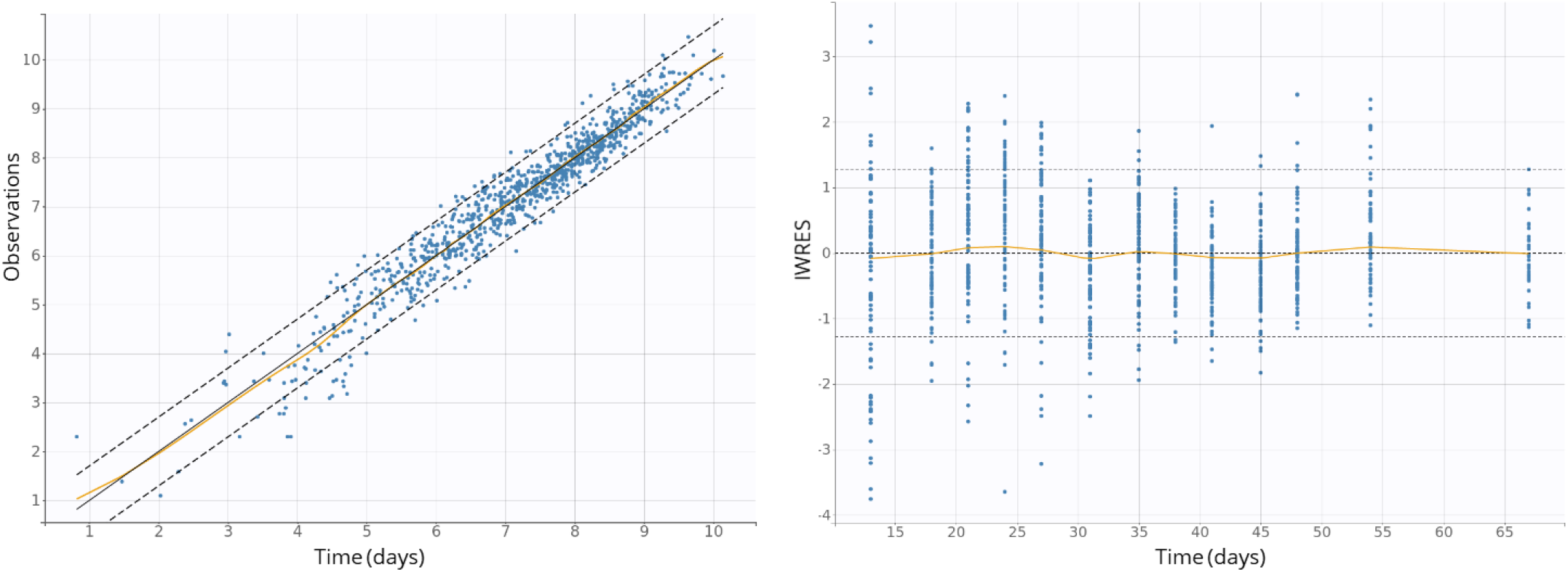
Standard Goodness-of-Fit Diagnostic Plots. On the left is individual predictions vs. observations and on the right are the individualized weighted residuals (IWRES) vs time. During model fitting, observations were natural-log-transformed to stabilize predictions. Therefore, residuals, predictions, and observations are natural-log-transformed in these figures. The predictions are approximately normally distributed. On the left, the one-to-one prediction line is the center solid black line, the spline (average agreement between individual prediction and observation) is solid orange, the dashed black lines are the borders of the 90% prediction interval. On the right, the zero residual error line is the center dashed black line, the spline is solid orange. The dashed black lines are the borders of the 90% prediction interval.

**Table 1.** Pharmacodynamic Model Parameters for Tumor Proliferation in NSCLC-Xenoαrafted Mice. Model parameter estimates (fixed and random-effects) as well as standard errors as determined by the Stochastic Approximation Expectation-Maximization algorithm as implemented in Monolix 2018R2.

| Model Parameter | Estimate | SE | RSE (%) | IIV | IIV (CV%) |
| --- | --- | --- | --- | --- | --- |
| $\alpha$ | 0.77 day<sup>-1</sup> | 0.0081 | 1.06 | 0.037 | 4.76 |
| $\beta$ | 0.04 day<sup>-1</sup> | 0.0005 | 1.16 | 0.015 | 33.55 |
| $\gamma$ | 35.74 (mg·day)<sup>-1</sup> | 6.36 | 17.79 | 0.46 | 1.28 |
| $\delta$ | 3.73 (mg·day)<sup>-1</sup> | 0.92 | 24.70 | 0.35 | 9.51 |
| $\tau_{\text{bound}}$ | .119 days | 0.0017 | 1.50 | 0.017 | 14.50 |
| $V_0$ | 3.4 RFU* | 0.27 | 22.55 | 1.73 | 143.03 |
| $k$ | 0.30 days | - | - | - | - |
| Residual Error Variance | 0.43 | 0.01 | 2.55 | - | - |

\*RFU: Relative Fluorescence Unit

### Simulations

Simulating the experimental treatments with a range of administration gaps from 0 to 10 days (step-size = 0.1 day) suggested that the optimal time delay between scheduling bevacizumab and PEM/CIS in mice is 2.0 days (**Simulation set 1**).

The simulated IIV of the optimal gap was relatively small. Only three values of *individual* optimal gap were produced. 96.5% of the virtual animals had an *individual* optimal gap of 2.0 days, 1. 0% of the virtual animals had an *individual* optimal gap of 2.1 days, and 2.5% of virtual animals had an *individual* gap of 1.9 days (**Simulation set 2**).

Scaling the dosage of bevacizumab to either 30 mg/kg or 10 mg/kg produced no effect in the estimated optimal gap and produced no effect in the IIV of the optimal gap (**Simulation set 3**).

Simulations of the typical human response to chemotherapy and bevacizumab were performed using IV administration, two-compartment absorption, and first order elimination models and parameters(32–34). Dosage and frequency of administration recommendations for BEV-PEM/CIS were adapted from DailyMed, a product label database maintained by the U.S. National Library of Medicine(35). Average adult weight and BSA were obtained from Center for Disease Control and literature estimates respectively(36,37).

Except for the proliferation rate of the tumor cells and rate of exponential decrease of the tumor relative growth rate (i.e. *α* and *β*), the PD model and parameterization were reused exactly as they were determined in the mouse portion of the model. *α* and *β* estimates were obtained from Bilous et al. 2018(38), where clinical NSCLC doubling times reported in Friberg and Mattson, 1998(39) were used to estimate population *α* and *β* for NSCLC in humans. The value of V_c_ came from the classical assumption that a 1 mm^3^ volume of tumor cells is approximately 10^6^ cells(40). V_0_ was arbitrarily set to 3 cm^3^.

The full PK/PD model was then used to simulate the typical cancer growth under various administration schedules with a starting tumor volume of 3 cm^3^. Parameter estimates are reported in **Table 2** and simulation summaries are depicted in **Figure 5**. The estimated optimal gap between bevacizumab and PEM/CIS administration in humans was 1.2 days.

**Figure 3.**
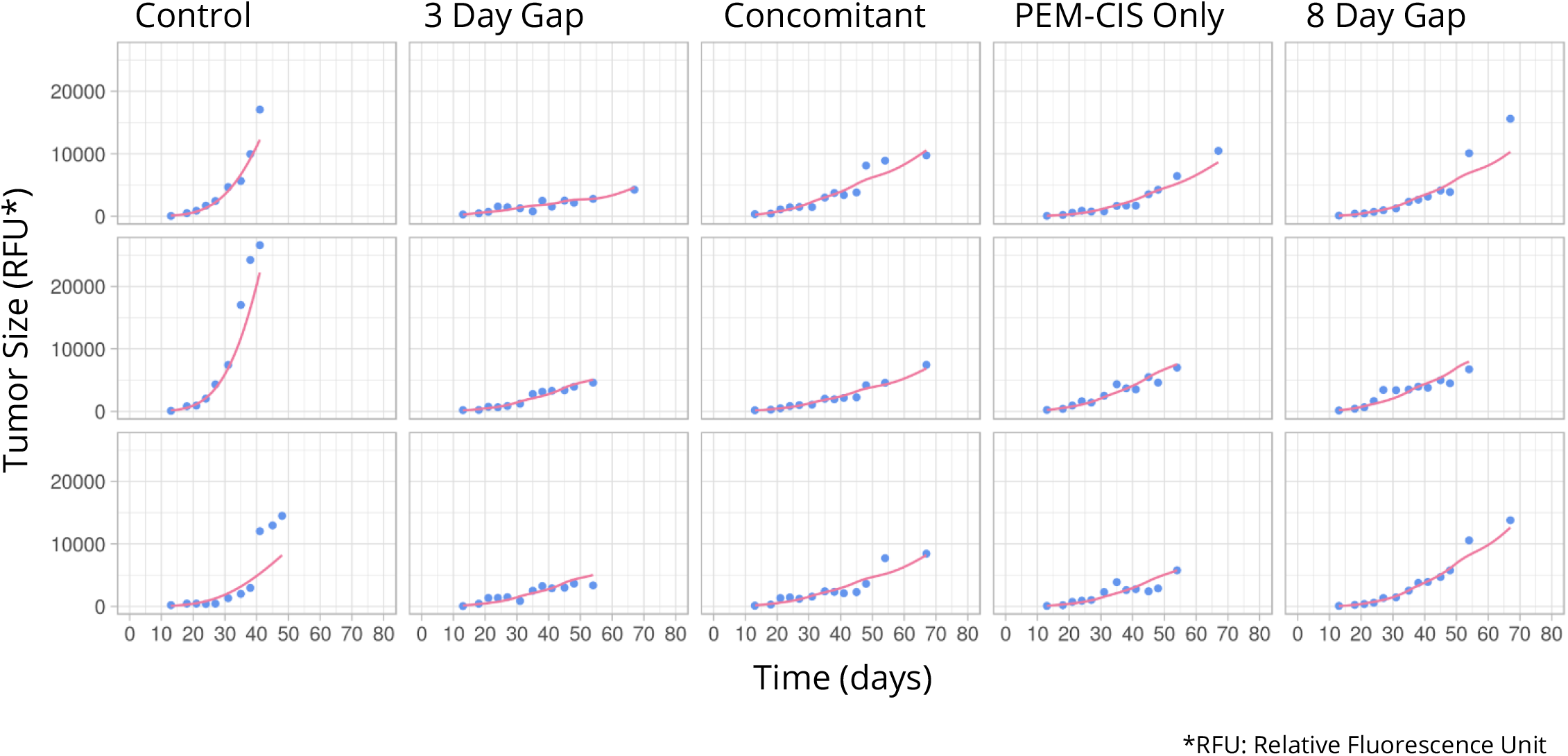
Sample of Individual Fits. The blue dots represent individual observations while the solid violet line represents individual fits.

**Figure 4.**
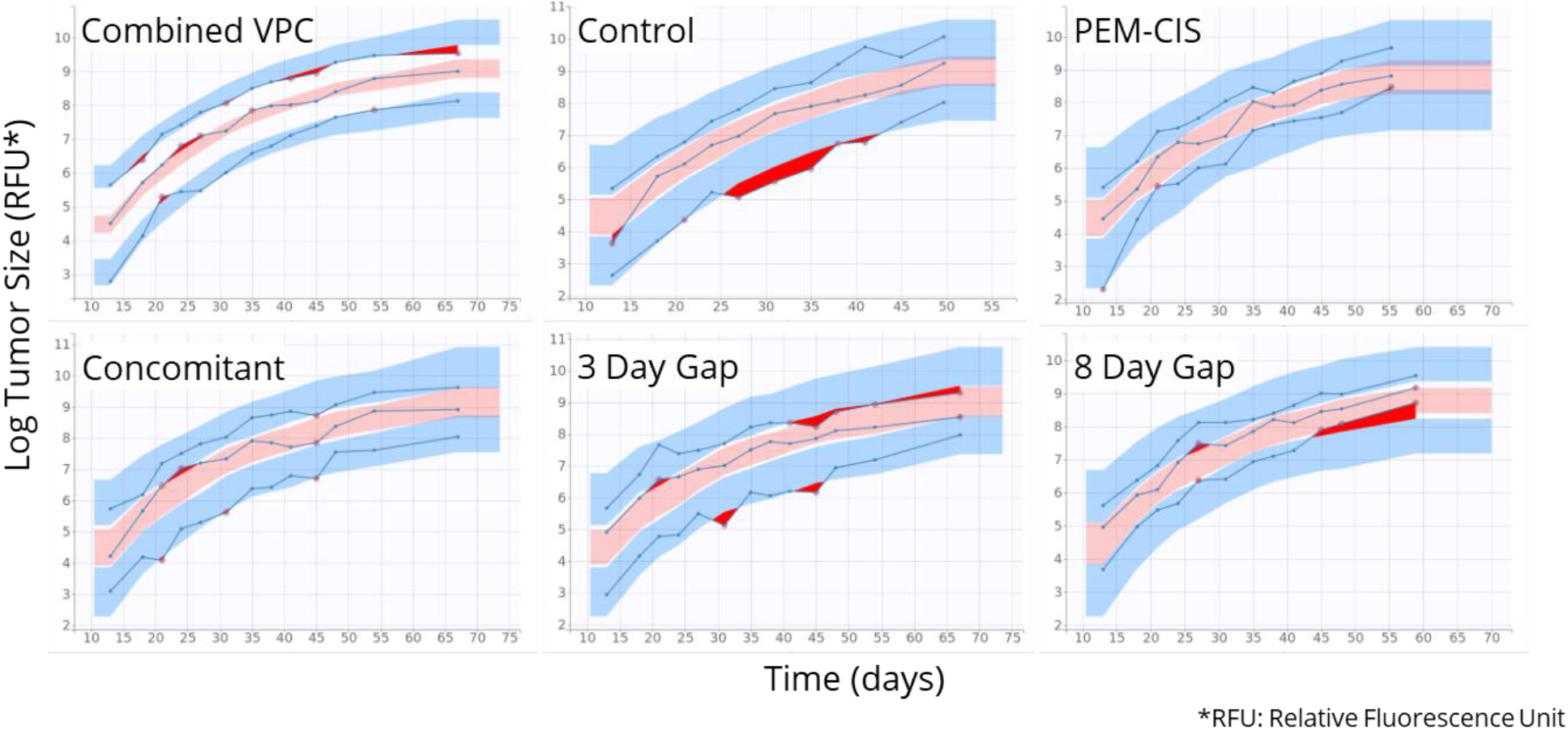
Combined and Stratified Visual Predictive Checks. The blue lines are the 10th, 50th, and 90th empirical percentiles calculated for each unique value of time. Blue and pink areas represent 90% prediction intervals for the 10th (blue), 50th (pink), and 90th (blue) percentiles. Prediction intervals are calculated by Monte Carlo simulation. To create prediction intervals for each unique value of time, 500 simulations are performed using random individual parameters. The red areas and red-circled points represent areas where empirical measurements fall outside of the bounds of the 90% orediction intervals.

**Figure 5.**
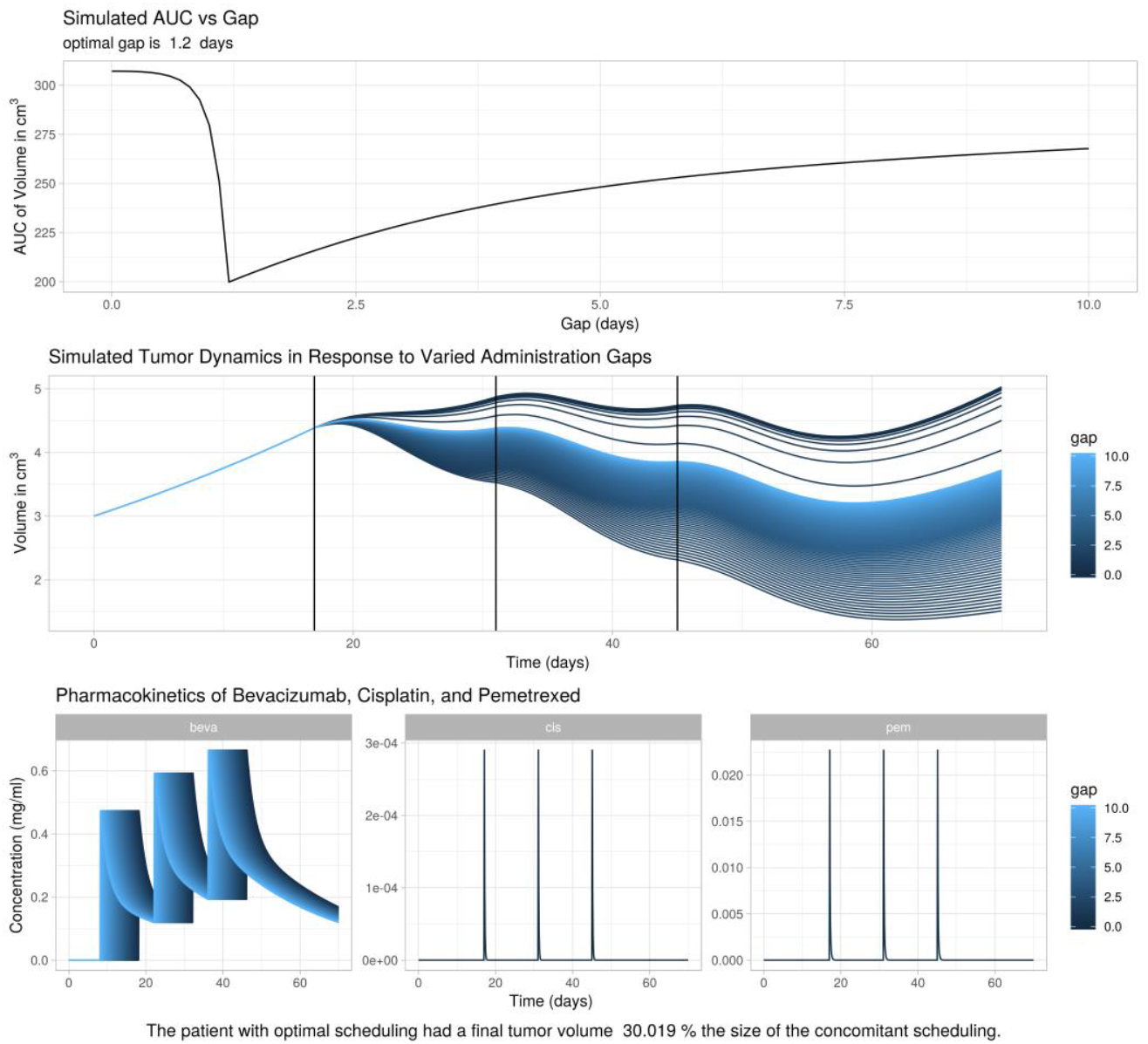
Human Pharmacodynamic and Pharmacodynamic Simulations Summary. To produce these figures, bevacizumab (BEV) was administered anywhere from 0 to 10 days (in steps of 0.1) before pemetrexed/cisplatin (PEM/CIS) was administered. Tumor growth was simulated from 0 to 67 days with no IIV and no RSE. In the top figure, AUC of tumor growth vs gap (0 to 10 days) is depicted. In the middle figure, tumor dynamics over time, with gap indicated by color, are depicted. In the bottom panel, the PK of BEV-PEM/CIS is depicted with gap indicated by color. The top figure indicates that the optimal scheduling gap is 1.2 days and the middle figure depicts the difference in tumor volume between administration gaps. The patient with optimal scheduling had a final tumor volume approx. 30% the size of the concomitant scheduling.

**Table 2.** Model Parameterization for Simulations of NSCLC treated with BEV-PEM/CIS in Humans. Except for the proliferation rate of the tumor cells and rate of exponential decrease of the tumor relative growth rate (i.e. *α,β*), the PD model and parameterization were reused exactly as they were determined in the mouse portion of the model. *α* and *β* estimates were obtained from Bilous et al. 2018(38), where clinical NSCLC doubling times reported in Friberg and Mattson, 1998(39) were used to estimate population *a* and *β* for NSCLC in humans. The value of *V_c_* came from the classical assumption that a 1 mm^3^ volume of tumor cells is approximately 106 cells(40). *V*_0_ was arbitrarily set to 3 cm^3^. *V, k_m,n_*, and *Q* represent volume, compartmental clearance from compartment m to compartment n, and intercompartmental clearance respectively.

| Pemetrexed Pharmacokinetics |  | Bevacizumab Pharmacokinetics |  | Pharmacodynamics |  |
| --- | --- | --- | --- | --- | --- |
| Clearance | 91.6 mL/min | Clearance | 0.22 L/day | $\alpha$ | 0.0284 day<sup>-1</sup> |
| $V_1$ | 12.9 L | $V_1$ | 2.8 L | $\beta$ | 1.03E-3 day<sup>-1</sup> |
| $Q$ | 14.4 mL/min | $k_{12}$ | 0.22 day<sup>-1</sup> | $\gamma$ | 35.7 (mg·day)<sup>-1</sup> |
| $V_2$ | 3.38 L | $k_{21}$ | 0.22 day<sup>-1</sup> | $\delta$ | 3.73 (mg·day)<sup>-1</sup> |
| Cisplatin Pharmacokinetics | | | | $\tau$ | 1.19 days |
| Clearance | 37.0 L/h | | | $V_0$ | 3 cm<sup>3</sup> |
| $V_1$ | 22.7 L | | | $k$ | 0.3 days |
| $Q$ | 10.7 L/h | | | $V_c$ | 10E-3 mm<sup>3</sup>·cell<sup>-1</sup> |
| $V_2$ | 19.5 L | | | | |

## Discussion

By normalizing tumor vasculature, bevacizumab improves delivery of PEM/CIS to tumors which increases PEM/CIS anti-tumor efficacy. Pemetrexed and cisplatin each have a narrow therapeutic window, low clearance, and high toxicity. It is therefore critical that BEV-PEM/CIS doses are administered as efficiently as possible. This makes BEV-PEM/CIS a natural fit for modeling and simulation studies, as the drug scheduling can be optimized without the need for multiple time and resource intensive *in vivo* studies. In this analysis we conducted an *in silico* study of the optimal administration of BEV-PEM/CIS in a xenograft, and human, model of NSCLC by constructing a mathematical model of tumor dynamics in response to BEV-PEM/CIS. In constructing that model, we were able to validate and refine previous modeling in BEV-PEM/CIS. Greater precision in parameter estimates was achieved through external estimation of *V*_c_, external validation of the residual error model, as well as using the larger fluorescence dataset to obtain final parameter estimates. Then, after exploring a range of predictions in mice, we scaled our model to predict optimal scheduling in humans.

The molecular profile of xenografts has a high degree of similarity with the molecular profile of the primary tumors from which the xenograft was derived(41) This indicates that there is compositional heterogeneity between xenografts and primary tumors. In addition, human NSCLC H460 cells are an experimentally established paradigm for modeling NSCLC tumors(42–44). Xenografts were grown after subcutaneous implantation in the mice flank, and not directly in the lungs. This is because monitoring tumor growth of lung orthotopic xenografts with non-invasive techniques is limitingly difficult due to both the rapid movements of the chest in mice during imaging, and photon emission attenuation by the high amount of water in lung tissues. Consequently, ectopic xenografts were considered as the more robust experimental model for data collection. Taken together, these considerations motivated our choice of experimental model and are the basis for which we scaled our mathematical model for making predictions in humans.

In the error modeling portion of the experiment, we demonstrated a strategy through which the choice of the error model can be validated externally to the primary dataset by including supplementary data collection in the experimental design. This simplified the error modeling step in the model building process.

The next stage of this study consisted of determining whether the semi-mechanistic model of tumor dynamics in response to BEV-PEM/CIS developed in Imbs et al. 2017(17) best fit the unfit fluorescence data from the same study. During model building, we attempted to balance our model building procedure between model performance (empirical fit) and the underlying biology, an approach often referred to as the *middle-out* approach(45,46).

In selecting potential PD models of tumor growth, several semi-mechanistic models were fit to the experimental data. The Gompertz model and linear-exponential model performed comparably. The parameters of the Gompertz model were estimated with greater precision than the parameters of the linear-exponential model (RSE) where the linear-exponential model was fit with a lower BIC than the Gompertz model. Ultimately, the Gompertz model was chosen over the linear exponential model due to the physiological relevance of its construction.

The parameterization of the final model was slightly unstable due to modest overparameterization. To compensate for this, *k* was fixed to a reasonable physiological estimate to improve precision of parameter estimates, and the search for *τ* was upper bounded to reduce spurious individual parameter estimates. In addition, the direct anti-proliferative effect of bevacizumab, and individual effects of pemetrexed and cisplatin, respectively, were removed from the model.

The modeling phase of the study resulted in several validations of previous findings. First, we confirmed the validity of the mathematical model previously published in Imbs et al. 2017(17), which we fit to our dataset. In doing so, we reconfirmed the ability of the model to describe BEV-PEM/CIS scheduling. We also reconfirmed the efficacy improvement of BEV-PEM/CIS dosing over PEM/CIS or control. We observed that a 3 day gap in scheduling is superior to both concomitant scheduling and an 8 day gap in scheduling. We were also able to build on previous work by identifying with greater precision the parameters underlying the mathematical model of BEV-PEM/CIS in NSCLC-tumor bearing mice.

In our mouse simulations, the final tumor volume (after 67 days) in the optimal scheduling group with BEV-PEM/CIS (gap = 2.0 days) was 88.5% of the size of final tumor volume in the concomitant scheduling group. This is consistent with our experimental results i.e. that mice administered bevacizumab approximately 2 days before PEM/CIS have a moderately better response (i.e. greater tumor size reduction) to BEV-PEM/CIS than mice who are administered BEV-PEM/CIS concomitantly.

The small magnitude of improvement in 2.0 day gap over concomitant scheduling is likely due to sub-optimal dosage and frequency of administration of BEV-PEM/CIS in the mice. We observed, by exploration, that a more robust preclinical response might be achieved by doubling the frequency of doses in mice and increasing the individual dosages by 50%.

We also found, through simulation, that scaling the dose of bevacizumab had no effect on the optimal gap and that IIV on gap is low.

Predictions made by our model agree with previous findings in BEV-PEM/CIS scheduling. The order of the optimal scheduling delay (2.0 days) is within the 1 to 5 day gap predictions of previous studies(12,18,19). Studies in tumor perfusion and bevacizumab showed Day 1 and Day 4 decrease in tumor perfusion which is consistent with the marginal predictions in our study i. e. optimal perfusion should be on the order of 2 days with comparable marginal losses on either side of that minimum(47).

After exploring various predictions in mice made by the model, the PK portion of the model was re-parameterized and the parameters of the PD portion of the model were scaled to simulate the relationship between varied administration schedules (i.e. gap) and efficacy in humans. Using this adapted parameterization, we estimated both optimal schedule of administration of BEV-PEM/CIS in humans and the marginal effects of a sub-optimal administration schedule of BEV-PEM/CIS in humans (i.e. bevacizumab administered too many or too few days before PEM/CIS).

In our human simulations, we predicted a robust improvement in response to sequential BEV-PEM/CIS relative to concomitant scheduling. The final tumor volume (after 67 days) in the optimal scheduling group (gap = 1.2 days) was 30% of the size of the final tumor volume in the concomitant scheduling group. If these predictions are accurate, scheduling optimization could result in significant improvement in BEV-PEM/CIS CT efficacy with no increase in toxicity.

The predicted scale of the increase in efficacy after scheduling optimization was much greater in the human simulations than was either predicted by the mouse simulations, or was measured empirically in mice. This is at least partially due to the greater study that has been undertaken in optimizing dosages for human treatment of NSCLC. However, these promising estimates should not be substituted for clinical testing.

When exploring marginal efficacy loss in sub-optimal administration schedules, we consistently found that the marginal cost of scheduling bevacizumab and PEM/CIS too close together in time was greater than the marginal cost of scheduling bevacizumab and PEM/CIS with too great of a gap in administration – in both mice and humans. This indicates that any potential clinical studies in antiangiogenics and cytotoxics should weight scheduling recommendations toward scheduling at slightly too large of gap.

Finally, the tumor microenvironment is known to be complex and varied. Tumor tissues contain necrotic pockets, heterogenous and dynamic microvasculature, and various sub-mutations which result in differential local growth rate and drug sensitivity. Considering this biological heterogeneity would greatly improve future model predictions and scalability between species.

In summary, our analysis confirms previous findings in BEV-PEM/CIS scheduling while improving precision of parameter estimates, improving prediction quality and detail, and scaling the model to predict the optimal scheduling of BEV-PEM/CIS in humans.

Antiangiogenics will continue to be useful agents in oncology. There are currently several other antiangiogenics regularly used in combination with cytotoxics which could potentially benefit from sequential administration (i.e. antiangiogenic then cytotoxic)(48). Of note, bevacizumab is currently only approved for concomitant administration with chemotherapy in all of its indications e.g. lung cancer, breast cancer, gastric cancer, etc. This contrasts with the optimized sequential scheduling that model simulations suggest.

There is a recent trend to develop model-informed drug development to optimize anticancer therapy. Our work highlights how mathematical modeling could help to refine clinical treatment modalities. The semi-mechanistic nature of this model allows it to be modularly reconfigured to extend predictions to other antiangiogenics as well as novel therapeutic paradigms such as the immune checkpoint inhibitor, monoclonal antibody pembrolizumab(49). This work continues to lay the foundation for building systems pharmacology models of the effect of antiangiogenic and antiproliferative combination therapy in advanced NSCLC. Tortuous vasculature is a phenotype exhibited by many solid tumors, and predicting optimal antiangiogenic scheduling could greatly increase the efficacy of future oncology therapeutics and combination therapies(50).

## Acknowledgements

We would like to thank Yeon-Jung Seo of Iowa State University College of Veterinary Medicine, Ames, IA, U.S.A, for performing the quality control assessment on code written to produce this study.

We thank Dr Valerie Le Morvan and Professor Jacques Robert from the Institut Bergonié, France, for their kind assistance.

## Author Contributions

B.K.S., S.B., and J.M. wrote the article. J.C., F.B., and S.B. designed the previous research. A.B. and J.C. performed the previous research. B.K.S., S.B., and J.M. analyzed the data.

## Abbreviations

adm: Administer, administered, administration, etc.
BIC: Bayesian Information Criteria
BSA: Body Surface Area
CT: Combination Therapy
DNA: Deoxyribonucleic acid
Gap: Short for the administration gap between (first) bevacizumab and (second) pemetrexed/cisplatin
IP: Intraperitoneal
IIV: Inter-individual variability
IV: Intravenous
NSCLC: Non-Small Cell Lung Carcinoma
PD: Pharmacodynamic
PK/PD: Pharmacokinetic/pharmacodynamic
PEM/CIS: Pemetrexed/cisplatin
BEV-PEM/CIS: Pemetrexed/cisplatin-bevacizumab
PK: Pharmacokinetics
ROS: Reactive Oxygen Species
RFU: Relative Fluorescence Units
sGOF: Standard Goodness-Of-Fit
VPC: Visual Predictive Check

