## Supplementary Figures for "Optimal Scheduling of Bevacizumab and Pemetrexed/cisplatin Dosing in Non-Small Cell Lung Cancer"

**Table S1** Administration (adm) schedule for the five experimental groups. The cytotoxics consisted of an administration of both 100 mg/kg of pemetrexed IP and 3 mg/kg of cisplatin IP. Bevacizumab was administered in doses of 20 mg/kg IP.

| Group | Experimental Day of bevacizumab adm | Experimental Day of cytotoxics adm | Group Size |
|---|---|---|---|
| Control | - | - | 15 |
| Cytotoxic control | - | 14, 28, 42 | 15 |
| No gap in adm | 14, 28, 42 | 14, 28, 42 | 16 |
| Three day gap | 11, 25, 39 | 14, 28, 42 | 15 |
| Eight day gap | 6, 20, 34 | 14, 28, 42 | 15 |

**Table S2** Administration (adm) schedule for the simulated human individual. Drugs were administered on three separate occasions, the 17th, 31st, and 45th experimental day. Bevacizumab administration (adm) was varied between simulation run i.e. it was administered anywhere between 0 and 10 days before Pemetrexed-Cisplatin.

| | Bevacizumab | Pemetrexed | Cisplatin |
|---|---|---|---|
| Adm Day 1<br>Experimental Day | 17 - gap | 17 | 17 |
| Adm Day 2<br>Experimental Day | 31 - gap | 31 | 31 |
| Adm Day 3<br>Experimental Day | 45 - gap | 45 | 45 |
| Dose<br>Given IV | 15 mg/kg | 100 mg/m$^2$ of BSA | 500 mg/m$^2$ of BSA |
| Duration of Infusion | 1 hours | 7 hours | 2 hours |

**Figure S1** In this figure the various error models are compared. In the left column are the probability density distributions (red) vs. theoretical normal distributions (blue) for each error group. On the right are the quantile-quantile plots which compare the quantiles between the theoretical and observed distributions of points. Below each set of sGOF graphics are the scores for the Kolomogorov-Smirnov normality test as well as the Pearson chi-square normality test. Graphically, the log-constant error model outperformed the constant error model. Numerically, the log-constant error model outperformed the constant error model in both the Kolmogorov-Smirnov normality test and the Pearson-Chi square test.

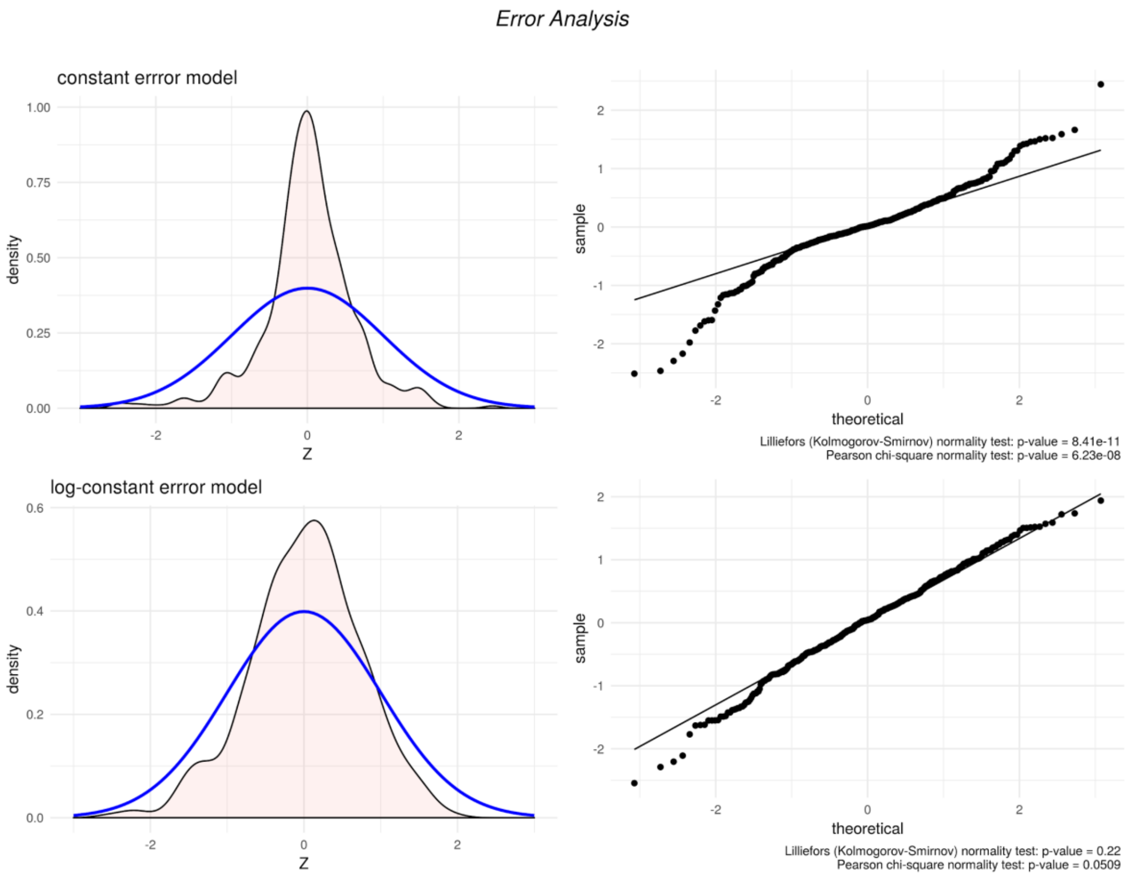

**Figure S2** Correlation between parameter estimates. The Pearson's Correlation Coefficient is displayed in each correlation and the red line is the linear regression of the conditional distribution.

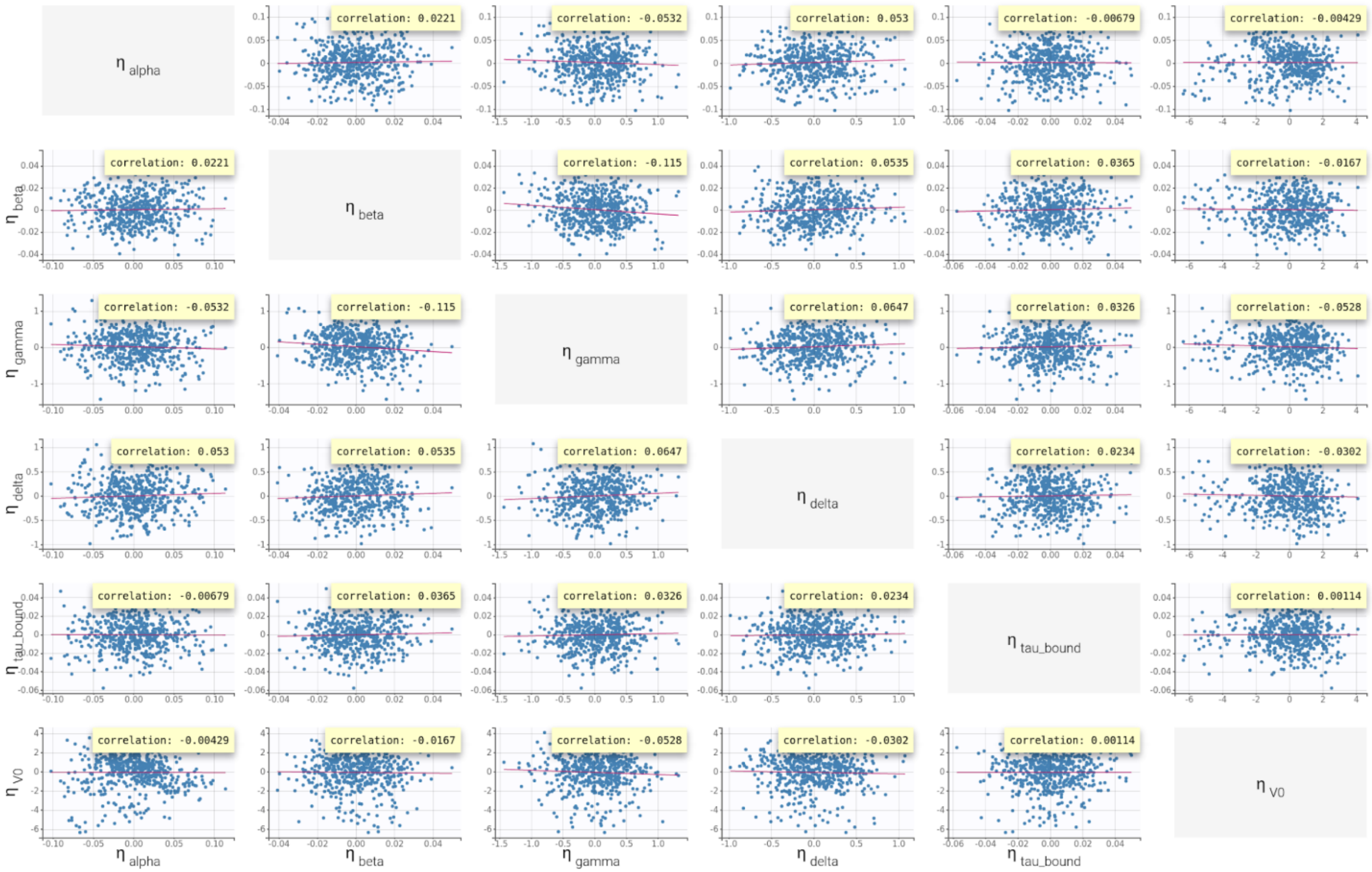

**Figure S3** Below are various SAEM convergence results produced by Monolix's *assessment suite*. The best performing (defined as lowest -2 *loglikelihood*) parameterization (in yellow) closely matches our final parameter estimates. The assessment indicates that there is some degree of instability in parameter estimates. But, variance in parameter estimates is relatively low and RSE on final parameter estimates is low, indicating that we've achieved a reasonable balance between the empirical fit and underlying biology. Especially important to our analysis is the relatively low variability in Tau parameter estimates.

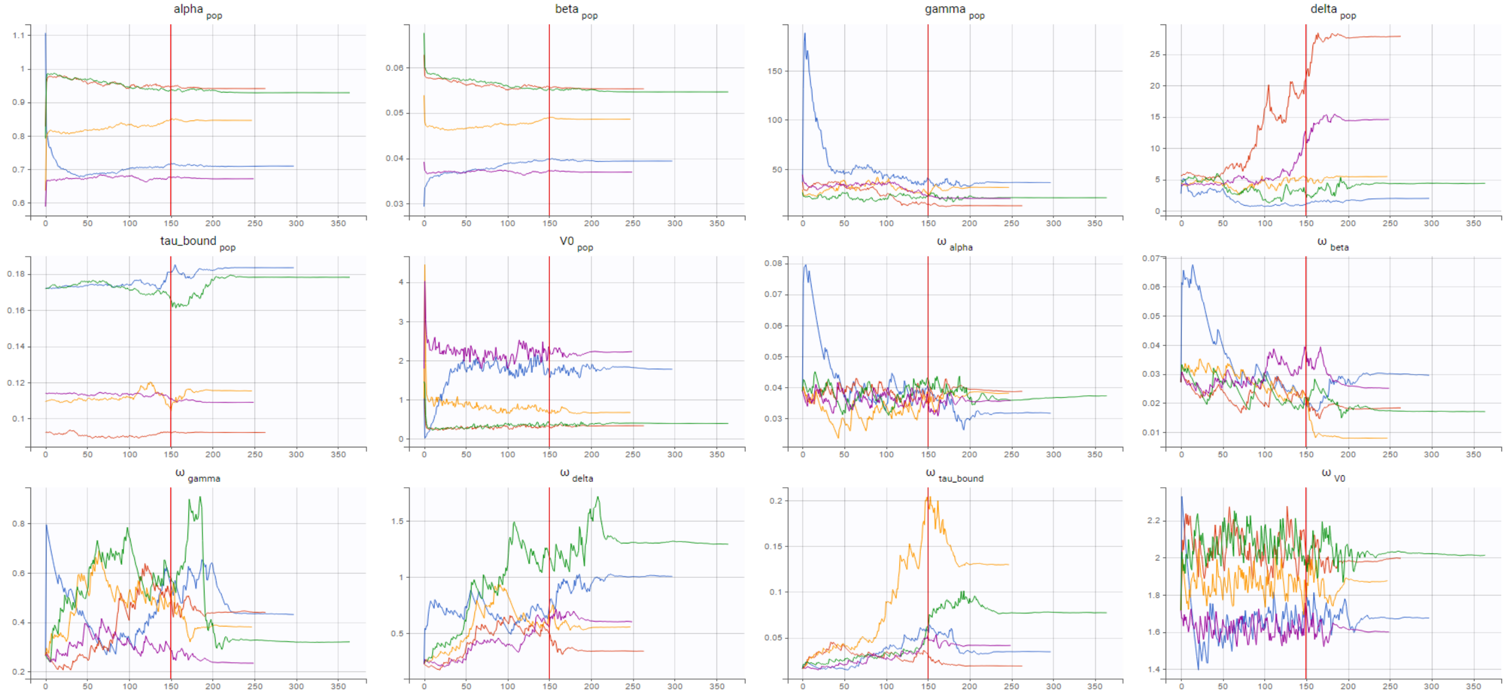

**Figure S4** Below is the 3D surface of simulated tumor growth response (dimension 1) with respect to gap (dimension 2) and time (dimension 3) in mice. This 3D surface allowed us to more intuitively understand the effect of varied administration schedules. It is available for manipulation at https://plot.ly/~Benjamin-PKPD/15/#/.

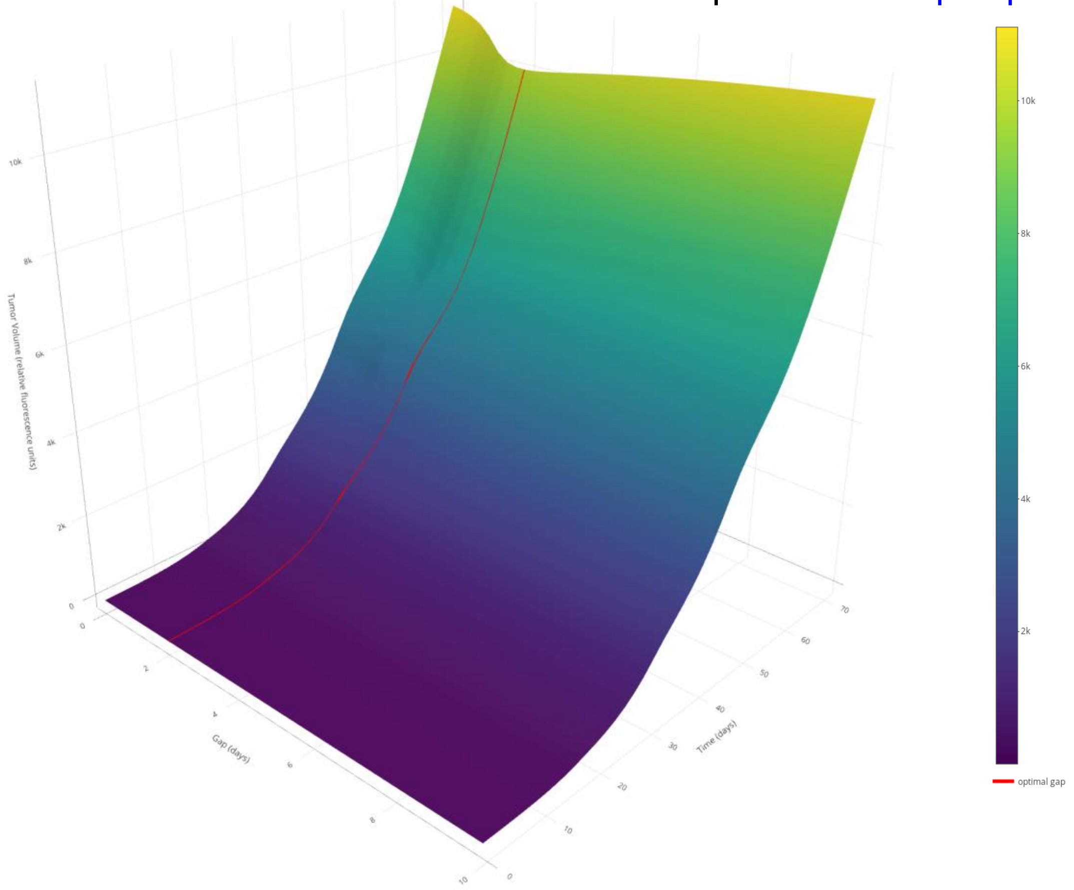

**Figure S5** Below is the 3D surface of simulated tumor growth response (dimension 1) with respect to gap (dimension 2) and time (dimension 3) in humans. This 3D surface allowed us to more intuitively understand the effect of varied administration schedules. It is available for manipulation at https://plot.ly/~Benjamin-PKPD/17/#/.

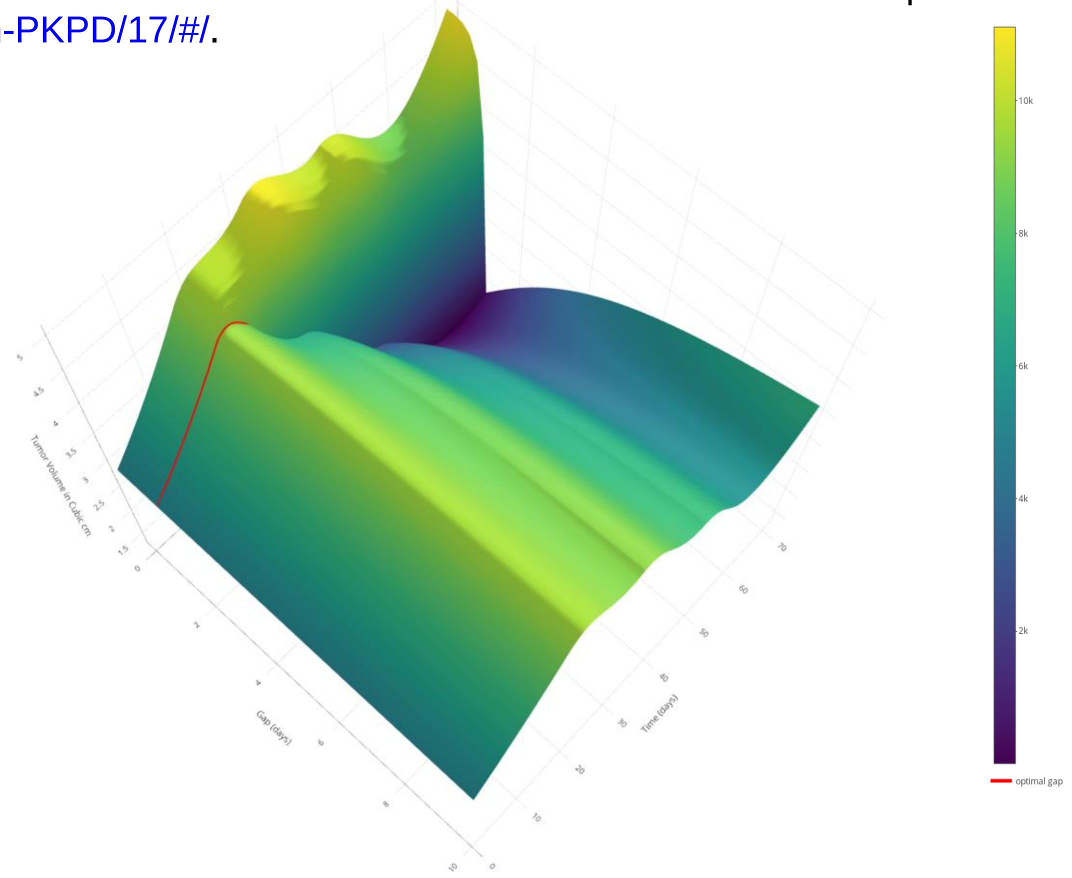
