## Supplementary Methods for "Optimal Scheduling of Bevacizumab and Pemetrexed/cisplatin Dosing in Non-Small Cell Lung Cancer"

**Supplementary Methods: Simulation Details**

In **simulation set 1**, population response was simulated with no IIV and without RSE. The simulated treatment schedule consisted of 100 mg/kg of pemetrexed IP and 3 mg/kg of cisplatin IP on Day 14, 28, and 42. 20 mg/kg IP of bevacizumab was virtually administered anywhere from 0 to 10 days (in steps of .1) before Day 14, 28, and 42. See **Table S2** for details.

The simulation was stopped after 67 virtual days. These simulations allowed the prediction of the optimal gap between administering bevacizumab and PEM-CIS. Efficacy of the drugs was assessed using AUC of the tumor growth over time as calculated by the spline method implemented in the R package DescTools 0.99.25[1]. The optimal gap was defined as the administration scheduling gap which resulted in the lowest AUC of tumor growth vs. time over the simulated time period (t = 0 to t = 67).

In **simulation set 2** the inter-individual variability of optimal gap was meticulously calculated using a virtual population of 1000 simulated mice. First, a set of parameters representing 1000 virtual mice was generated without RSE from the parameter distributions determined in the structural modeling phase. Then, the procedure from **simulation set 1** was replicated per individual to estimate each *individual* optimal gap. Finally, the distribution of *individual* optimal gaps was assessed using the mode, standard deviation, and range as well as graphical methods such as a violin plot of count density.

To produce **simulation set 3**, the procedure which generated **simulation set 1** was repeated with scaled doses of bevacizumab. First, 10 mg/kg IP of bevacizumab was substituted for 20 mg/kg IP of bevacizumab. Second, 30 mg/kg IP of bevacizumab substituted for 20 mg/kg IP of bevacizumab. This was done to assess the effect of bevacizumab dosage on the estimated optimal gap.

In **simulation set 4** an analogous treatment regimen to **simulation set 1** was simulated in a virtual NSCLC tumor-bearing human population using adapted human dosing recommendations. The random effects were set to zero and population response was simulated

---

[1]Aho ASIR source code and/or documentation previously published by (in alphabetical order): K, Alfons A, Anderegg N, et al. DescTools: Tools for Descriptive Statistics.; 2019. https://CRAN.R-project.org/package=DescTools. Accessed January 23, 2019.

without RSE. Therefore, our simulations represent the average effect of varied administration schedule.

The model used in **simulation set 4** was a scaling of the model fit obtained from the xenograft mouse model. The PK portion of the model was built and parameterized using previously published PK models and typical parameter values. The PD parameter estimates were obtained from both the xenograft mouse model and from literature values.

Tumor dynamics in response to PEM-CIS-BEV were simulated using a scheduling gap of 0 days to 10 days in steps of .1. A graphical representation of the AUC vs. administration gap was generated to determine the optimal gap between bevacizumab and PEM-CIS dosing.
